# Repair of leukemia-associated single nucleotide variants via interallelic gene conversion

**DOI:** 10.1101/2024.04.16.587991

**Authors:** Alexander J. Silver, Donovan J. Brown, Sarah D. Olmstead, Jackson M. Watke, Agnieszka E. Gorska, Londa Tanner, Haley E. Ramsey, Michael R. Savona

## Abstract

CRISPR-Cas9 is a useful tool for inserting precise genetic alterations through homology-directed repair (HDR), although current methods rely on provision of an exogenous repair template. Here, we tested the possibility of repairing heterozygous single nucleotide variants (SNVs) using the cell’s own wild-type allele rather than an exogenous template. Using high-fidelity Cas9 to perform allele-specific CRISPR across multiple human leukemia cell lines as well as in primary hematopoietic cells from patients with leukemia, we find high levels of reversion to wild-type in the absence of exogenous template. Moreover, we demonstrate that bulk treatment to revert a truncating mutation in *ASXL1* using CRISPR-mediated interallelic gene conversion (IGC) is sufficient to prolong survival in a human cell line-derived xenograft model (median survival 33 days vs 27.5 days; *p* = 0.0040). These results indicate that IGC can be applied to numerous types of leukemia and can meaningfully alter cellular phenotypes at scale. Because our method targets single-base mutations, rather than larger variants targeted by IGC in prior studies, it greatly expands the pool of risk-increasing genetic lesions which could potentially be targeted by IGC. This technique may reduce cost and complexity for experiments modeling phenotypic consequences of SNVs. The principles of SNV-specific IGC demonstrated in this proof-of-concept study could be applied to investigate the phenotypic effects of targeted clonal reduction of leukemogenic SNV driver mutations.

## INTRODUCTION

A major challenge for anti-cancer therapies is to target neoplasia while sparing healthy tissue. Somatic mutations that arise with aging are known to contribute to disease pathogenesis or disease severity, and constitute one class of distinguishing features with the potential to be specifically targeted (1). Single nucleotide variants (SNVs) are highly represented in sequenced cancer genomes (2). In the blood, there are numerous recurrent age-associated acquired SNVs which have been identified in both clonal hematopoiesis (CH), a pre-disease state, and myeloid malignancy (3,4). Because each SNV represents only a slight deviation from the wild-type sequence, there are inherent challenges with ensuring proposed modalities exploiting cancer-associated SNVs maintain a high degree of specificity (5). On the other hand, the small footprint of SNV lesions makes them attractive targets for genetic manipulation via homology-directed repair (HDR), as the success rate of HDR has an inverse relationship with the size of the alteration to be made (6). While HDR is traditionally carried out by providing an exogenous repair template, it should also be possible to use HDR to replace a heterozygous SNV if a double-stranded break in DNA can be induced on the mutant allele while leaving the wild-type allele intact to act as a template. To this end, sporadic cell-intrinsic HDR following the targeting of a heterozygous SNV by CRISPR knockout (CRISPR KO) has been observed in zygotes (7-9). Many cancer-associated SNVs are heterozygous, but so far there exist no studies examining whether interallelic gene conversion (IGC) aimed at missense/nonsense SNVs could intentionally be used to repair deleterious point mutations in somatic human cells.

In this study, we report that acquired SNVs that arise in CH and are associated with hematologic malignancy can be efficiently restored to their wild-type sequences using widely available CRISPR reagents without the need for exogenous HDR templates. We show that use of high-fidelity Cas9 allows for the specific targeting of mutant alleles. We find that incorporation of the SNV into a standard 20-basepair gRNA protospacer sequence minimizes off-target cutting of the wild-type allele. Furthermore, we demonstrate that use of a commercially available HDR enhancer can drastically reduce insertions/deletions (indels) generated during this process. In sum, these experiments show that efficient reversion of heterozygous gain-of-function or loss-of-function SNVs in human leukemia cell lines and primary patient samples can be accomplished using routine CRISPR workflows. The simplicity of this approach should enable easy adoption into any number of research applications, especially among those in the scientific community who have experience performing CRISPR KO experiments. This method, which in its simplest form needs only high-fidelity Cas9 and gRNA, is a resource-efficient approach to studying SNV-mediated cellular pathologies, such as high-risk CH and mechanisms of leukemogenesis. However, it has the potential to be broadly applied to research questions in basic and medical translational science, especially in the field of oncology.

## MATERIALS & METHODS

### Ethical statement

Experiments were conducted on deidentified primary patient samples collected and distributed by the Vanderbilt-Ingram Cancer Center Hematopoietic Malignancies Repository following acquisition of written informed consent, and in accordance with the tenets of the Declaration of Helsinki and approved by the Vanderbilt University Medical Center Institutional Review Board (#151710).

### Cell culture

K562 cells (ATCC, Manassas, VA, USA) were cultured in RPMI 1640 (Corning, Corning, NY) supplemented with 20% fetal bovine serum (FBS; R&D Systems, Minneapolis, MN, USA) and 1% penicillin/streptomycin (P/S; Thermo Fisher Scientific, Waltham, MA, USA). OCI-AML3 (DSMZ, Germany) cells were cultured in MEM α (Thermo Fisher Scientific) with 20% FBS and 1% P/S. OCI-AML5 (DSMZ) cells were cultured in MEM α supplemented with 20% heat-inactivated FBS, 1% P/S, and 10 ng/mL of GM-CSF (PeproTech, Rocky Hill, NJ, USA). THP-1 cells (ATCC) were cultured in RPMI 1640 with 10% FBS, 1% P/S, and 0.05 mM β-mercaptoethanol (MilliporeSigma). SET-2 (DSMZ) cells were cultured in RPMI 1640 with 20% FBS and 1% P/S. Leukapheresis samples were cultured in RPMI 1640 with 5% FBS, 0.1 mM β-mercaptoethanol, and 10 ng/mL IL-1β (PeproTech), 10 ng/mL IL-3 (PeproTech), and 10 ng/mL GM-CSF.

### CRISPR/Cas9

Using a Neon transfection system (Thermo Fisher Scientific), cells were electroporated with HiFi Cas9 (IDT, Newark, NJ, USA) at 150 μg/mL complexed with sgRNA (IDT) at a 1:2.5 ratio. For electroporation, cell lines were resuspended in Buffer R, while primary cells were resuspended in Buffer T, all at ∼100 x 10^6^ cells/mL. Settings were as follows: K562, 1450 V, 10 ms pulse width, 3 pulses; OCI-AML3, 1600 V, 10 ms pulse width, 3 pulses; OCI-AML5, 1500 V, 30 ms pulse width, 1 pulse; THP-1, 1700 V, 20 ms pulse width, 1 pulse; SET-2, 1400 V, 20 ms pulse width, 2 pulses; primary cells, 1650V, 10ms pulse width, 3 pulses. For experiments including HDR Enhancer V2 (IDT), cells were resuspended in media containing 1 μM of the compound for overnight culture, followed by media exchange the following morning. Genomic DNA was isolated using DNA Blood Mini Kit (Qiagen, Hilden, Germany) at 48 hours post-electroporation unless otherwise indicated.

### Measurement of post-CRISPR allele fractions

Sanger sequencing of PCR-amplified CRISPR loci was performed through Genewiz/Azenta using the primers listed in Supplementary Table S1. Primers were purchased through IDT. Sanger trace files were visualized with SnapGene Viewer.

Next-generation sequencing was performed at the VUMC VANTAGE core using the VUMC Clonal Hematopoiesis Sequencing Assay v2.0, a custom capture protocol which covers 22 frequently mutated CH genes, including the full exonic sequences of all of the genes examined in this study, plus several dozen germline SNPs (10). Samples were sequenced to a depth of between 0.5 x 10^6^ to 1 x 10^6^ reads per sample on a NovaSeq sequencer (PE150). The DRAGEN Somatic pipeline (v3.10.4, Illumina) was used to validate SNVs and generate BAM files mapped to GRCh38/hg38 that were then analyzed in R (v4.1.1) with *Rsamtools* (v2.10.0). Prior to final quantification of allele frequency and indels at the targeted SNV, reads were filtered to include only those with unique mapping (MAPQ ≥ 60) and coverage of both the SNV and at least four bases to either side of the predicted cut site 3 bp upstream of the PAM, in order to reduce the likelihood of underestimating indel proportions. Data were then reviewed manually in IGV (v2.14.0, Broad Institute). Allele fractions for non-targeted heterozygous SNPs were derived from IGV coverage statistics for reads with MAPQ ≥ 60. Ideograms were generated with *RIdeogram* (v0.2.2) mapping to GRCh38/hg38.

### Time course experiment

K562 and OCI-AML5 cells from three independent CRISPR experiments per cell line were cultured continuously for six weeks with regular splitting. Genomic DNA was isolated at the beginning of the experiment (day 0) and every subsequent week. Isolated DNA was submitted for NGS sequencing as above in a single batch and VAFs were analyzed as described above. Timepoints with very low sequencing coverage (< 100 reads at SNP of interest) were excluded from the analysis to minimize the possibility of spurious results from imprecise estimates.

### Animal model

Animal experiments were performed in accordance with guidelines approved by the Institutional Animal Care and Use Committee (IACUC) at Vanderbilt University Medical Center. Experiments used female NOD.Cg-*Prkdc^scid^ Il2rg^tm1Wjl^* Tg(CMV-IL3, CSF2, KITLG)1Eav/MloySzJ (NSGS) mice (The Jackson Laboratory #013062) (11), aged 6 to 8 weeks old, which were irradiated with 1 Gy radiation. All experiments used female mice. Animals were randomized to receive cells from the treatment or control conditions. Each mouse received 5 x 10^6^ K562 cells that had been electroporated with high-fidelity Cas9 and either scramble or SNV-specific gRNA. Cells were transplanted immediately following electroporation. Two independent experiments were conducted and the results pooled. Survival analysis was performed in R using *survival* (v3.5.7), *survminer* (v0.4.9), and *ggsurvfit* (v1.0.0).

### Western blot

Protein lysates were extracted from 1-5 million cells using RIPA buffer (Thermo Fisher Scientific) supplemented with cOmplete Protease Inhibitor Cocktail (MilliporeSigma, Burlington, MA, USA) and phosphatase inhibitor cocktail set V (Calbiochem, San Diego, CA, USA). Lysates were mixed 1:1 with 2X Laemmli Buffer (Bio-Rad, Hercules, CA, USA) supplemented with β-mercaptoethanol and denatured at 95°C for 10 minutes. Samples were run on 4-20% TGX gels (Bio-Rad) at 100V for 5 min, then 80V for 2 additional hours. Wet transfer to PVDF membrane (MilliporeSigma) was performed at 100V for 60 min on ice using transfer buffer prepared in house (20% v/v methanol, 0.3% w/v Tris, 1.44% w/v glycine). After overnight blocking in 5% milk in TBST, membranes were probed with primary antibody for 60 min at room temperature, followed by 3xTBST washes, then probed with goat anti-rabbit secondary antibody at 1:5000 for 45 minutes at room temperature, followed by 3xTBST washes. Chemiluminescent detection using autoradiography film (Thomas Scientific, Swedesboro, NJ, USA) was performed using SuperSignal West Dura (Thermo Fisher Scientific) for ASXL1 and SuperSignal West Pico PLUS (Thermo Fisher Scientific) for β-actin. The primary antibodies used in this study were polyclonal anti-ASXL1 raised in rabbit (PA5-68360, Thermo Fisher Scientific) and polyclonal anti-β-actin raised in rabbit (A2066, MilliporeSigma).

### RNAseq

Two million cells from each condition (scramble or Y591X-targeted gRNA) were harvested from three independent CRISPR experiments on K562 cells. RNA was extracted using the RNeasy Mini Kit (Qiagen), treated with DNAse I (NEB, Ipswich, MA, USA) and subsequently purified with the Monarch RNA Cleanup Kit (10 μg) (NEB). Library preparation using NEBNext® Poly(A) selection and sequencing were performed at the VUMC sequencing core. Samples were sequenced to a depth of 50 million reads on a NovaSeq 6000 sequencer, PE150. Preprocessing was performed on the DRAGEN RNA Pipeline (v3.6.3). Raw counts were normalized using *DESeq2* (v1.34.0). Differential expression analysis was conducted with *DESeq2*Enrichment analysis, and plotting was performed using *clusterProfiler* (v4.10.0) and *enrichplot* (v1.22.0).

### Cell number and proliferation assays

For cell expansion experiments, cells were plated at an initial density of 100,000 cells/mL and cultured for 72 hours. Counting of cells stained with trypan blue 0.4% (Thermo Fisher Scientific) was performed every 24 hours using a Countess II FL automated cell counter (Thermo Fisher Scientific). For cell division quantification, cells were incubated in CellTrace Violet (Thermo Fisher Scientific) in PBS for 20 minutes at room temperature in the dark, followed by three washes in PBS. Immediately following, an aliquot of cells was fixed in 4% paraformaldehyde, while the remaining cells were cultured for 72 hours followed by paraformaldehyde fixation. Samples were analyzed on a 3-laser BD LSRFortessa (BD Biosciences, Franklin Lakes, NJ, USA). Cell division bins were determined by taking the mean day 0 fluorescence, dividing sequentially by 2, and then calculating the midpoints between the halved intensity values. For cell cycle analysis, cells were plated at an initial density of 200,000 cells/mL. After 24 hours in culture, 7.5 x 10^5^ cells were washed twice in PBS, then fixed by the dropwise addition of 3 mL cold 70% ethanol. Cells were then incubated at -20°C for 60 minutes, washed three times in PBS, and resuspended in 0.5 mL BD Perm/Wash Buffer (BD Biosciences). Following this, 0.1 mL of cell suspension was incubated with human APC-conjugated Ki-67 antibody (Cat. 350514, BioLegend, San Diego, CA, USA) at room temperature in the dark for 30 minutes. Cells were washed twice and resuspended in 0.2 mL BD Perm/Wash Buffer. Samples were incubated with 5 µL Propidium Iodide Staining Solution (Cat. 556463, BD Biosciences) and analyzed on a 5-laser BD LSR II (BD Biosciences). Three independent experiments were performed for each assay. FACS data were analyzed using FlowJo software (v10).

### Visualization

Except as noted, plots were generated using *ggplot2* (v3.3.6), *ggpubr* (v0.4.0), and *RColorBrewer* (v.1.1.3).

### Statistical analyses

For analyses of variant allele fractions, protein quantification, cell division number, and cell cycle state two-sided Welch’s t-tests were used. Two-way ANOVA with replication was used to test for difference in cell line growth kinetics. To test for differential expression in the Beat AML cohort, a two-tailed Wilcoxon rank sum test was used. The log-rank test was used to analyze survival differences.

### Data availability

The RNAseq data used in this study are available in GEO under GSE212730. The cell line DNAseq data used in this study are available under NCBI SRA accession number PRJNA880841. The human subject DNAseq data are not deposited in a public repository due to patient privacy requirements but are available upon reasonable request.

## RESULTS

### SNV-directed CRISPR cutting enables interallelic gene conversion in leukemia models

Working under the hypothesis that IGC will occur in a situation where a heterozygous mutant allele is cut while the wild-type (WT) allele remains intact, we sought to test if a strategy using high-fidelity Cas9 and SNV-specific gRNAs without exogenous template could lead to correction of a mutant sequence. To test this, we looked across several human leukemia cell lines and identified different heterozygous malignancy-associated SNVs that could serve as potential targets. We then designed allele-specific gRNAs against these sequences, including the mutant SNV in the protospacer sequence (**Fig. 1A**). We then electroporated the parental cell lines with ribonucleic protein (RNP) consisting of the SNV-specific guide and a high-fidelity Cas9, followed by collection of DNA 48 hours post-electroporation (**Fig. 1B**). We observed both a relative increase in the measured amount of the WT allele fraction and a decrease in the mutant allele fraction as assessed by Sanger sequencing (**Fig. 1C**). To confirm this, we performed custom-capture next generation sequencing (NGS) with >300X coverage of target loci in order to assess the relative abundance of WT and mutant alleles. In five tested cell lines, we found a significant increase in the abundance of WT allele following treatment (**Fig. 1D-H**), accompanied by a relative decrease in the mutant allele fraction (Supplementary Fig. S1A-B). We also noted that this occurred even in cell lines harboring three copies of the targeted chromosome: THP-1 (2 mutant and 1 WT) (**Fig. 1F**) and K562 (1 mutant and 2 WT) (**Fig. 1H**). These results demonstrate that HDR can occur at heterozygous loci in cell lines even in the absence of exogenous repair templates.

**Figure 1.**
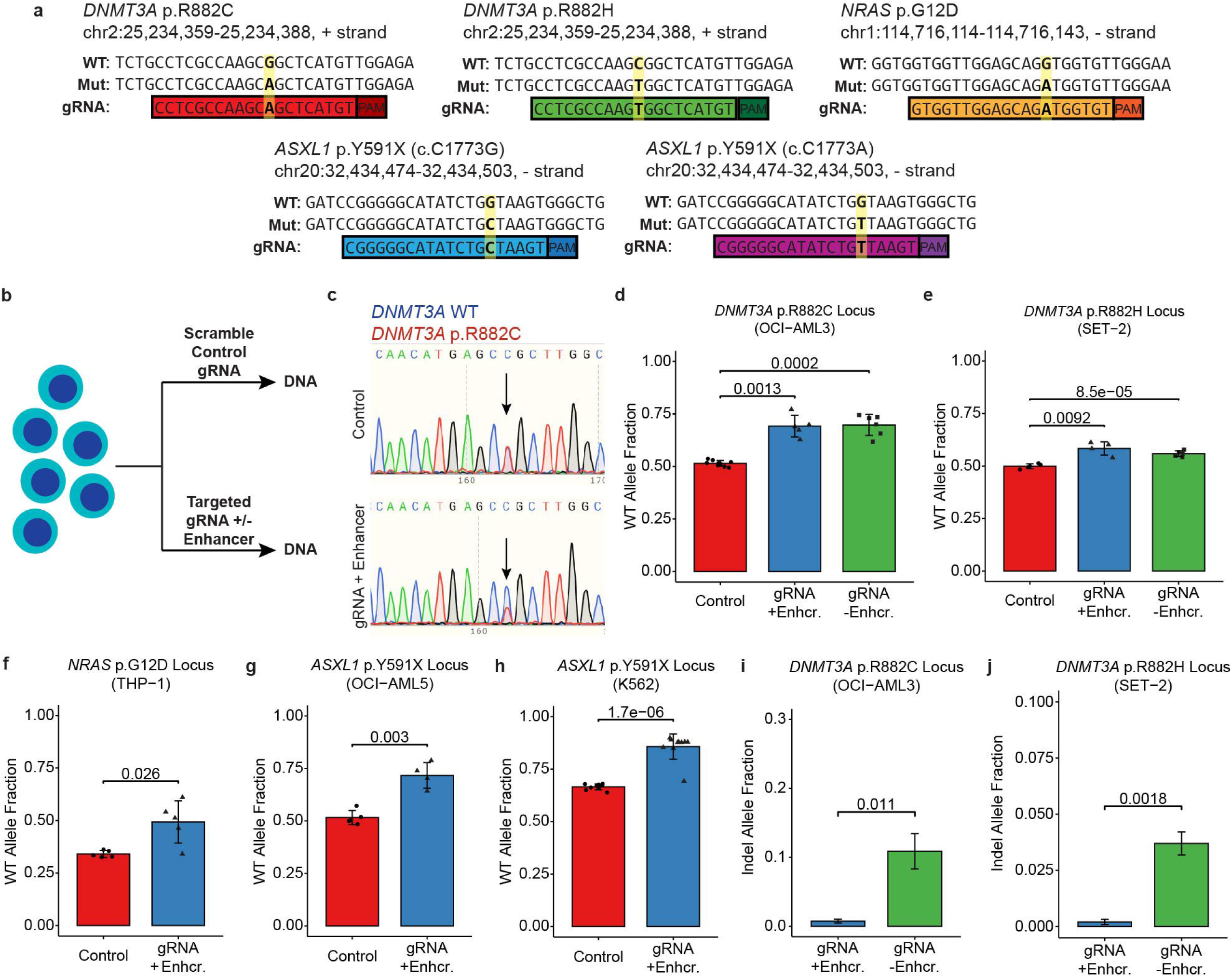
Interallelic gene conversion can increase wild-type allele fraction in hematopoietic cell lines. (A) The allele-specific gRNA sequences designed against *DNMT3A* p.R882C, *DNMT3A* p.R882H, *NRAS* p.G12D, *ASXL1* p.Y591X (c.C1773G), and *ASXL1* p.Y591X (c.C1773A). The position of the targeted mutant single nucleotide variant (SNV) is highlighted in yellow. Sequences are oriented such that the protospacer:PAM may be read left-to-right. (B) Cells were electroporated with ribonucleic protein complex containing SNV-specific or scramble gRNA with or without homology-directed repair (HDR) enhancer. (C) Sanger trace showing increase in the wild-type (WT) allele fraction and decrease in the mutant allele fraction of *DNMT3A* p.R882C in OCI-AML3 cells. The coding orientation is depicted, which is the reverse-complement of the locus shown in panel A. (D) The WT allelic fraction following control treatment (red), treatment with targeted gRNA plus HDR enhancer (blue), or gRNA without HDR enhancer (green) for the *DNMT3A* p.R882C locus in OCI-AML3 cells, (E) the *DNMT3A* p.R882H locus in SET-2 cells, (F) the *NRAS* p.G12D locus in THP-1 cells, (G) the *ASXL1* p.Y591X (c.C1773G) locus in OCI-AML5 cells, and (H) the *ASXL1* p.Y591X (c.C1773A) locus in K562 cells. (I) Total indels in samples receiving mutant-targeting gRNA plus HDR enhancer (blue) or gRNA without HDR enhancer (green) in OCI-AML3 cells and (J) SET-2 cells. Bar plots and error bars represent mean and standard deviation. P-values represent results from Welch’s two-sided t-test. See also Supplementary Fig. S1.

Because we primarily intended to test whether IGC could be performed at heterozygous SNVs at all and we therefore included an inhibitor of non-homologous end joining (“HDR enhancer”) in all of our experiments from the outset to achieve maximum biological effect, we also wished to test whether IGC would occur at meaningful levels or if indel formation would predominate in the absence of HDR enhancer. Thus, in the first two cell lines we tested, OCI-AML3 and SET-2, we performed experiments in the presence and absence of HDR enhancer. We found that the WT allele fraction was significantly increased even without HDR enhancer (**Fig. 1D-E**). We then phased the NGS reads at each locus to be able to quantify the overall and allele-specific burden of indels (**Table 1** and Supplementary Fig. S1C). This revealed that the total number of indels was significantly higher in the absence of HDR enhancer (**Fig. 1I-J**). Moreover, across all of our cell line experiments, only a small minority of reads contained indels affecting the WT allele (range 0.00%-0.91% of total reads; **Table 1**). This indicates that WT-sparing IGC can be accomplished using high-fidelity Cas9 and mutant-specific gRNAs with or without HDR enhancer; although, the use of enhancer minimizes the amount of indel formation in targeted cells.

**Table 1.** Allelic phase of indels in treated cell line samples.

| Sample | Target | HDR Enhancer? | Absolute Percent of Total Reads Containing Indels (SD) | Absolute Percent Total Reads: |  |  |
| --- | --- | --- | --- | --- | --- | --- |
|  |  |  |  | Indels Phased to WT Allele | Indels Phased to Mutant Allele | Indels Unable to be Phased |
| OCI-AML3 | *DNMT3A* p.R882C | No | 10.9% (6.3%) | 0.044% | 8.1% | 2.7% |
|  |  | Yes | 0.8% (0.6%) | 0.032% | 0.51% | 0.26% |
| SET-2 | *DNMT3A* p.R882H | No | 3.7% (1.1%) | 0.51% | 2.4% | 0.83% |
|  |  | Yes | 0.2% (0.2%) | 0% | 0.090% | 0.11% |
| THP-1 | *NRAS* p.G12D | Yes | 7.9% (7.6%) | 0.24% | 6.4% | 1.23% |
| OCI-AML5 | *ASXL1* p.Y591X | Yes | 15.2% (4.3%) | 0.91% | 3.2% | 11% |
| K562 | *ASXL1* p.Y591X | Yes | 8.5% (3.1%) | 0.76% | 1.25% | 6.5% |

### Longitudinal analysis demonstrates allele fraction changes are persistent and are not explained by chromosomal loss

While large-scale deletions and copy number changes have not been reported for allele-specific CRISPR in somatic cells, allele-specific editing in human embryos has been shown to lead to some chromosomal loss (12). This led us to investigate the possibility of whether large-scale genomic events were driving the change in allele fraction at the targeted SNV. To do this, we examined the allele fraction at all covered heterozygous germline SNPs on the same chromosome as the targeted SNV, reasoning that chromosomal loss events affecting the mutant SNV would also affect the allelic ratios of all heterozygous alleles *in cis* (**Fig. 2A-E**). We observed no indication of copy number alterations at the non-targeted heterozygous SNPs when we examined these data, with the exception of one K562 SNP. This SNP (Chr20:32,429,316) is located < 5 kb proximal to the targeted *ASXL1* SNV and had a small but statistically significant increase in allele fraction (**Fig. 2E**). A more distal K562 SNP did not have a significant change, which led us to postulate three possible explanations: 1) unresolved DNA repair in a fraction of cells, 2) IGC extending beyond the area immediately around the cut site, or 3) localized copy number alterations. Given that kilobase-magnitude IGC has been observed previously (7), we considered the former two hypotheses as the most likely; the location of *ASXL1* in close proximity to the centromere of chromosome 20 makes partial chromosomal copy number loss without a change in the allele frequency of the distal SNP seem the least plausible explanation.

**Figure 2.**
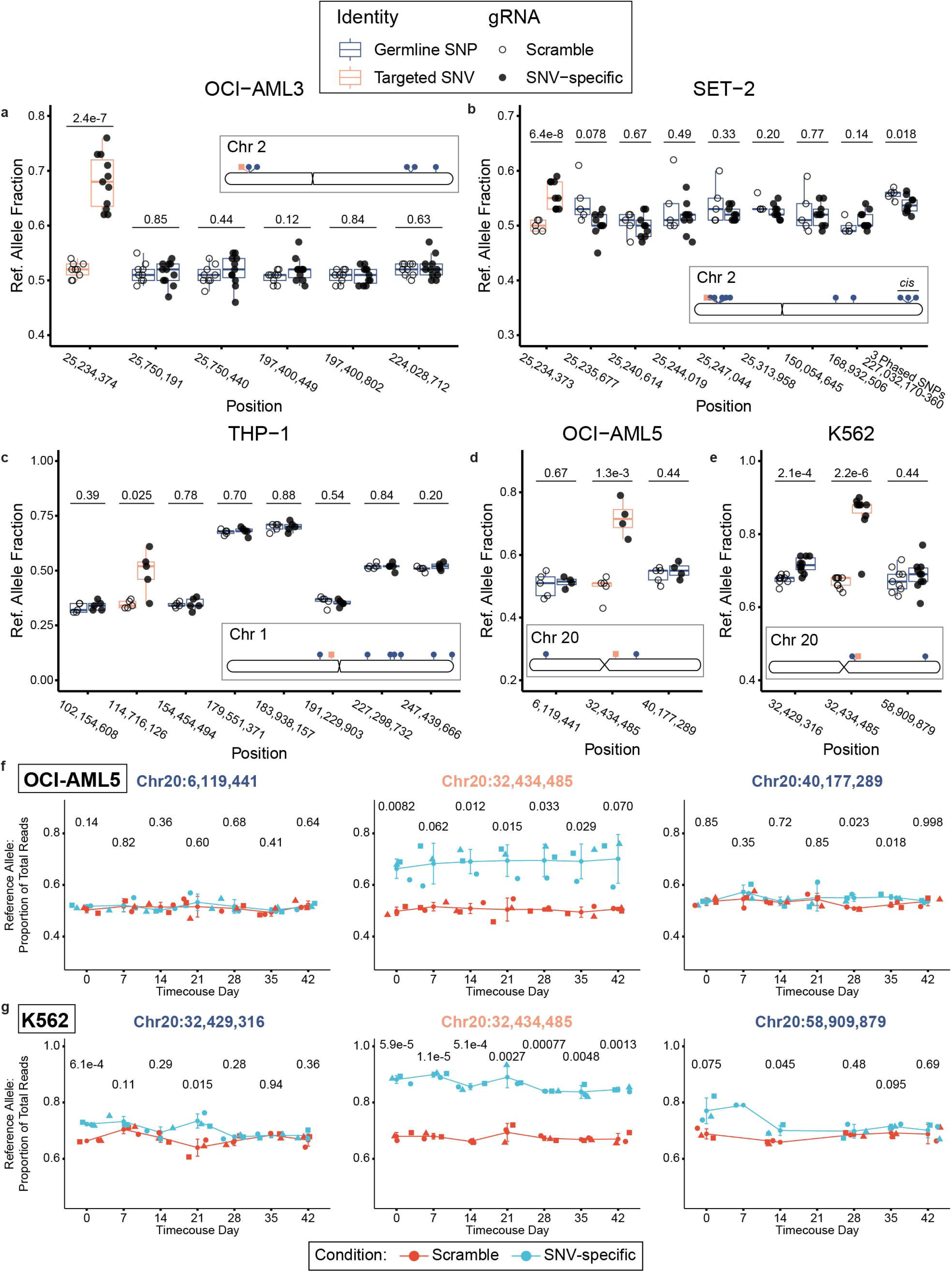
Allele fraction changes are persistent and are not explained by chromosomal loss. (A-E) The allelic fraction of the GRCh38/hg38 reference allele (Ref. Allele Fraction) for the targeted single nucleotide variant (SNV) and informative heterozygous germline single nucleotide polymorphisms (SNP) on the same chromosome for (A) OCI-AML3 cells, (B) SET-2 cells, (C) THP-1 cells, (D) OCI-AML5 cells, and (E) K562 cells. The inset ideograms show the relative positions of each of the depicted variants. Boxplots depict means and quartile ranges. (F-G) Time course data showing the reference allele fraction for targeted SNV and informative germline SNPs for (F) OCI-AML5 cells and (G) K562 cells. Experimental replicates are denoted with unique symbols. Mean and standard deviation are depicted. P-values represent results from Welch’s two-sided t-test.

In part to explore the cause of the altered allele fraction at this proximal SNP, we conducted a time course experiment with both K562 cells along with a cell line with both proximal and distal SNPs that lacked evidence of allelic shifts, OCI-AML5. In this experiment, we also wished to observe if the allelic ratios for targeted SNVs would change across time. To this end, we collected genomic DNA at weekly intervals over six weeks from these two cell lines. Next-generation sequencing of these samples revealed that, for both cell lines, the mean allele fraction at the targeted SNV and informative germline SNPs remained relatively constant across time (**Fig. 2F-G**). Moreover, the allele fractions for the SNV-specific gRNA treated *ASXL1* p.Y591X SNVs held steady for each experimental replicate. While the allele fraction for the K562 Chr20:32,429,316 SNP was significantly greater for the Y591X-specific gRNA arm at the earliest timepoint, it was more similar to scramble gRNA control at the following timepoints. This argues against IGC extending beyond the target cut site as well as localized copy number loss, both of which should engender persistent changes to allele fraction. Overall, these time series data indicate that the change to the allele fraction of the targeted SNV is a specific and stable event which is not explained by chromosomal loss, although copy number changes occurring in a small fraction of cells cannot be ruled out entirely.

### Bulk editing of *ASXL1* truncating mutation reduces cellular proliferation with concomitant transcriptomic changes

Having observed genetic evidence of IGC in several cell lines at various loci, we then wished to understand how bulk editing using this method might impact cellular phenotypes of genetically complex cancer cells. We focused on correction of a truncating mutation in *ASXL1*, due to this gene’s importance in myeloid cancers but also because correction of a stopgain SNV is a logical place to start in evaluating SNV-driven pathology: restoration of the stopgain SNV to a wild-type allele may lead to phenotypic differences, while introducing stopgain-causing frameshifts into an allele already harboring a stopgain mutation should be phenotypically indistinguishable. The ASXL1 protein is a part of several histone-modifying complexes, including polycomb repressive complex 2 (PRC2) and the BAP1 histone H2AK119Ub deubiquitinase (DUB) complex (13). Stopgain mutations in *ASXL1*, including p.Y591X, are commonly observed in clonal hematopoiesis (14,15) and myeloid malignancies (16), being associated with poor prognosis in the latter (17,18). We therefore examined the effects of *in vitro* IGC correction of *ASXL1* p.Y591X in K562 cells, which displayed robust conversion of mutant alleles to WT. When we treated K562 cells with *ASXL1* p.Y591X-targeting gRNA, we observed a significant reduction in the amount of mutant protein but no fluctuation in the amount of full-length ASXL1, as compared to controls (**Fig. 3A-C**). These results indicate that IGC of *ASXL1* stopgain SNVs preserves the expression of the WT protein and can lead to reduced levels of the mutant protein.

**Figure 3.**
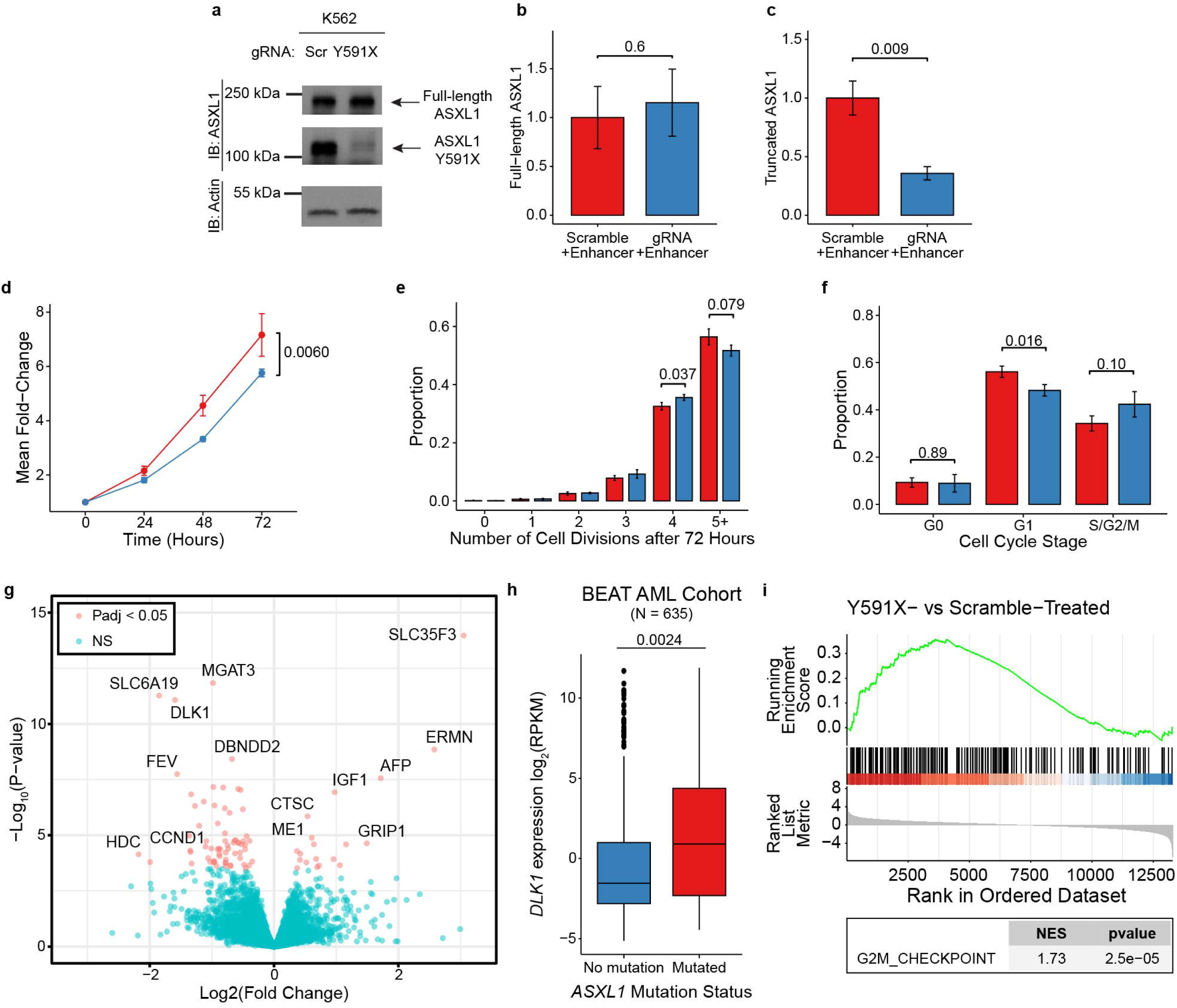
Reversion of *ASXL1* p.Y591X results in transcriptional downregulation of pro-growth pathways. (A) Western blot of scramble gRNA or Y591X-gRNA treated K562, showing full-length and truncated ASXL1 and β-actin. Protein quantification for (B) full-length ASXL1 and (C) truncated ASXL1 (N = 3 experimental replicates). (D) Fold expansion (mean +/- SEM) of cell count over 72 hours for K562 treated with scramble (red) or mutant-targeting (blue) gRNA (N = 3 experimental replicates). (E) Cell divisions (mean +/- SD) as measured by CellTrace Violet fluorescence of K562 treated with scramble (red) or mutant-targeting (blue) gRNA after 72 hours of growth (N = 3 experimental replicates). (F) Proportion of cells in G0, G1, and S/G2/M (mean +/- SD) as quantified by Ki-67 and propidium iodide fluorescence of K562 treated with scramble (red) or mutant-targeting (blue) gRNA (N = 3 experimental replicates). (G) Volcano plot showing RNAseq results of differentially expressed genes (FDR = 0.05) in Y591X-treated vs. scramble-treated conditions. (H) Transcriptional expression of *DLK1* in the Beat AML cohort, stratified by *ASXL1* mutation status. (I) Enrichment plot showing results from preranked gene set enrichment analysis for the Hallmark G2M Checkpoint pathway. P-values represent results from Welch’s two-sided t-test (B, C, E, F), two-way ANOVA with replication (D), and two-sided Wilcoxon rank sum test (H). See also Supplementary Fig. S2.

We then asked whether this bulk treatment would affect the cellular phenotype, particular cellular proliferation. We assessed K562 cell numbers over time for scramble or SNV-specific gRNA treated samples seeded at equal densities and saw a significantly lower fold expansion of the experimental samples (two-way ANOVA, p = 0.006; **Fig. 3D**). To determine if this decreased expansion was the result of a diminished rate of replication, we conducted a CellTrace assay and observed that the population of experimentally treated cells was shifted toward fewer divisions compared to controls (**Fig. 3E** and Supplementary Fig. S2A-B). Consistent with these findings, when we performed dual Ki-67/propidium iodide staining to look at stages of the cell cycle, we saw that while both conditions had a similar proportion of cells in G0, there were significantly fewer experimental cells in G1 phase, coupled with a relative increase in S/G2/M phases (**Fig. 3F** and Supplementary Fig. S2C). Altogether, these data suggest that IGC correction of the *ASXL1* p.Y591X SNV in K562s leads to a diminished proliferative drive and alters the cycling of these cells in a manner consistent with a delayed progression through cellular replication checkpoints.

We next turned to examine how the transcriptome was affected by the reduced burden of ASXL1 mutant protein. We performed bulk RNAseq on K562 cells treated with scramble gRNA or mutant-specific gRNA. One of the genes found to have significantly lower expression after IGC was *DLK1* (**Fig. 3G**), a gene frequently upregulated in myeloid malignancy (19). *DLK1* expression is known to be positively correlated with stemness/differentiation block in the hematopoietic niche (20) and has been shown to promote proliferation in a K562 model (19) and primary patient samples (21). An association between *ASXL1* mutation and *DLK1* expression has not previously been reported, so we were intrigued to find that *ASXL1* mutation does indeed have a significant association with higher *DLK1* transcription in the Beat AML cohort (**Fig. 3H**) (4,22,23). Additionally, consistent with our data suggesting slower progression through the cell cycle, we found that the Hallmark gene set (24) for the G2/M checkpoint was significantly enriched (**Fig. 3I**). Together, these data suggest the diminished proliferation of K562 cells following bulk editing of *ASXL1* p.Y591X via IGC may result from downregulation of pro-proliferative factors and partially retarded progression through cell cycle checkpoints.

### Correction of ASXL1 truncating mutations prolongs survival in myeloid malignancy xenograft model

We next sought to determine whether bulk-editing of cells to correct truncating mutations in *ASXL1* could improve survival in xenograft models. It has been previously reported that complete removal of truncated *ASXL1* in a subcloned KBM5 cell line delayed mortality in a mouse model (25), but it was unclear whether a survival benefit would be seen if instead the xenograft were comprised of a more heterogenous bulk-edited population in which some residual fraction of cells retain mutant protein, or even if the KBM5 results were generalizable beyond that particular model. We therefore transplanted K562 cells pre-treated with either scramble or *ASXL1* p.Y591X-targeting gRNA into sublethally irradiated NSGS mice (**Fig. 4A**). No significant difference was found in animal weights over time between conditions (**Fig. 4B**). We found the ASXL1-targeted group had significantly longer survival than the control group, with a median survival of 33 days compared to 27.5 days (*p* = 0.0040; N = 8 per arm) (**Fig. 4C**). These results indicate that the truncating *ASXL1* mutation contributes to the pathogenicity of K562 cells in the xenotransplant setting and that a single-time bulk CRISPR correction of this mutation is sufficient to impart a survival advantage in such a model.

**Figure 4.**
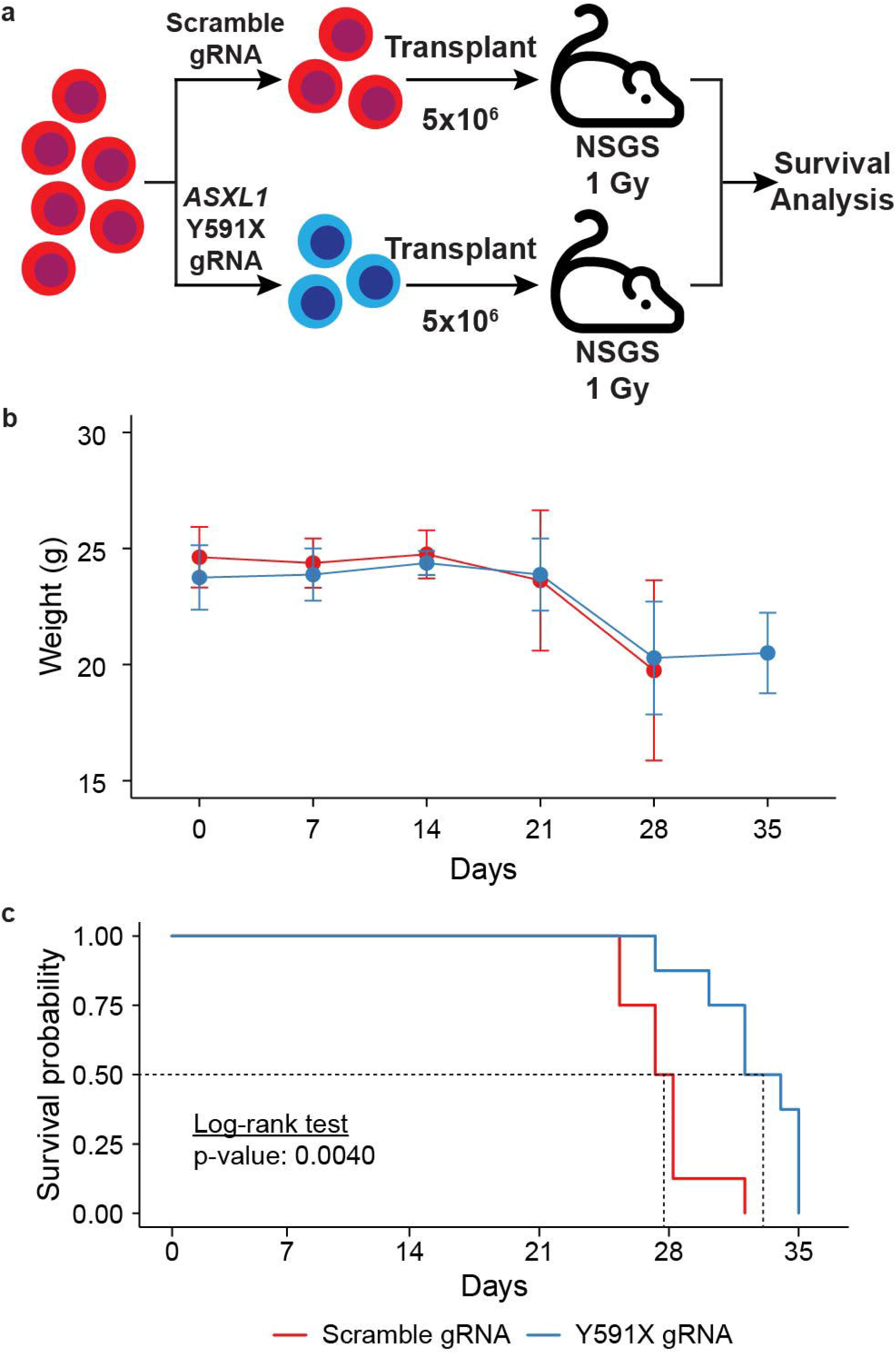
Bulk interallelic gene conversion is sufficient to prolong survival in cell-line derived mouse xenograft model. (A) Schematic of xenograft model. (B) Measured weights (mean +/- SD) and (C) survival curves of NSGS mice receiving human K562 AML cells treated with either scramble gRNA (red) or mutant-targeted gRNA (blue). Results based on two independent experiments (N = 8 per condition). Significance was determined using a log rank test.

### Primary hematopoietic cells are amenable to interallelic gene conversion

Having successfully tested IGC in multiple cell lines, we then asked whether it would be possible to perform invoke HDR without an exogenous repair template at a SNV in primary human cells. To examine this, we used previously frozen peripheral blood mononuclear cells (PBMCs) from two patients in AML blast crisis who underwent leukapheresis procedures, knowing that a large proportion of collected cells would be mutant blasts and thus suitable targets for our experiments. Clinical NGS analysis of peripheral blood samples done at the time of diagnosis indicated that AML 01 included an *ASXL1* p.Q748X mutation with an VAF of 42% while AML 02 harbored a *DNMT3A* p.R882H mutation measured at 44% VAF, both of which proved targetable by SNV-specific gRNAs (**Fig. 5A**). Following a single treatment with mutant-specific gRNA, we observed an increase in the WT allele fraction in both patient samples (**Fig. 5B-C**). At the same time, there were no statistically significant differences in the informative germline SNPs on the targeted chromosomes, indicating the VAF changes at the targeted SNV cannot be attributed to chromosomal loss. Notably, there were also very few total indels (≤ 8 total indels per replicate) measured in either patient sample following treatment despite high read coverage for the target locus (mean depth of 1153X for AML 01 and 1474X for AML 02). Although, both experiments were conducted using HDR enhancer with all replicates so it is impossible to determine whether there would be more indels in the absence of this drug (Supplementary Table S2). Taken together, these results indicate that SNV-specific IGC may also be accomplished in primary human hematopoietic cells from an AML disease context.

**Figure 5.**
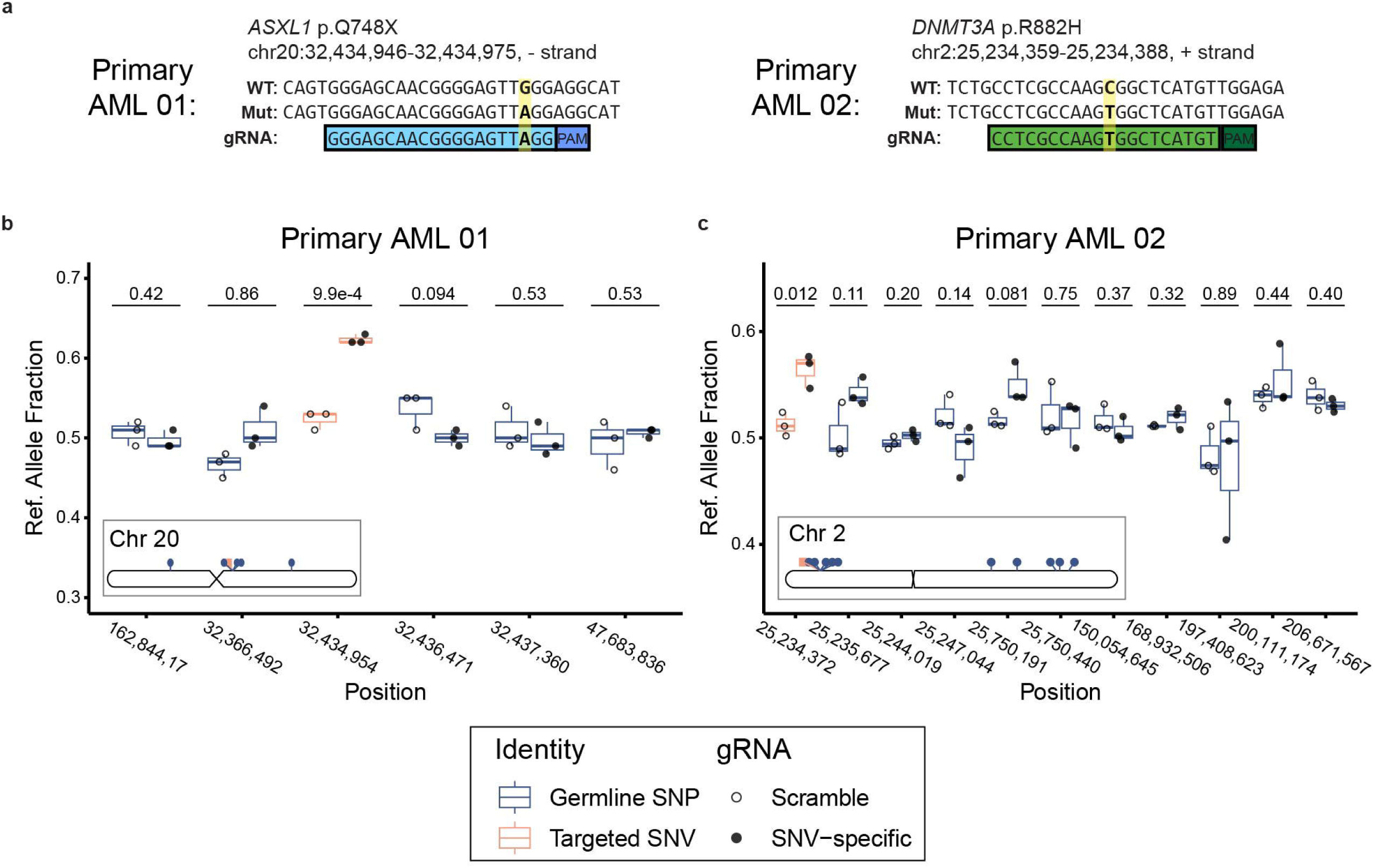
Successful interallelic gene conversion in primary leukemia cells. (A) The sequences of the *ASXL1* p.Q748X and *DNMT3A* p.R882H targeted gRNAs. Sequences are oriented such that the protospacer:PAM may be read left-to-right. (B-C) The allelic fraction of the GRCh38/hg38 reference allele (Ref. Allele Fraction) for the targeted single nucleotide variant (SNV) and informative heterozygous germline single nucleotide polymorphisms (SNP) on the same chromosome for (B) AML 01 and (C) AML 02. The inset ideograms show the relative positions of each of the depicted variants. Boxplots depict means and quartile ranges. P-values represent results from Welch’s two-sided t-test.

## DISCUSSION

In this study, we demonstrate that intentional IGC can be used to repair heterozygous leukemia-associated single-basepair mutations. We successfully tested an IGC approach on numerous SNVs across leukemia cell lines and primary patient samples and went on to show the utility of this approach as a preclinical research tool by targeting a truncating *ASXL1* mutation, which led to clear phenotypic alterations in the treated cells. Targeting *ASXL1* led to a partial amelioration of a proliferative cellular phenotype, and we identified a potential downstream target in *DLK1* that can be further investigated in future work. Finally, we show that a bulk IGC correction of a single SNV in a cancer cell line is sufficient to significantly prolong survival in a mouse xenograft model. The primary importance of these findings is that SNV-directed IGC is a straightforward and streamlined approach to revert mutant SNVs to their wild-type sequence in a large number of cancer-causing mutations previously not targeted in therapeutic or preventative fashion. This exponentially expands the potential targets for IGC-mediated repair since, to date, IGC has only been attempted on larger mutations or mutations affecting rare PAM sites.

Several additional points deserve emphasis. In our experiments using a high-fidelity Cas9 enzyme, the results indicate limited cutting of the wild-type allele, which we believe is crucial to the success of IGC. This is true despite our use of protospacer sequences that differ from the wild-type by only a single base (and ranging in position from 3-10 bp upstream of the PAM). We show that IGC can occur with RNP treatment alone, but that the mutant may be repaired with indels, an unintended outcome that can be minimized by using a small molecule inhibitor of nonhomologous end joining (NHEJ), which reduces indel formation by this error-prone DNA repair method and favors HDR. Lastly, we also find that IGC can be accomplished even when there is polyploidy for an allelic locus, a situation that is not uncommon in malignant cells (26).

There is much promise in the use of CRISPR-based methodologies to directly interrogate disease-causing mutations, and our work extends the basic research basis for this field. Previously described CRISPR HDR approaches have relied on exogenously provided repair templates, and this approach has led to new insights generated from a broad range of genetic knock-in models (27-31). The IGC method presented here offers an alternative to exogenous-template HDR in a highly circumscribed context: when the purpose is to replace a SNV existing in a cell with its alternate allele. In this arena, our method offers several potential benefits: 1) higher theoretical efficiency due to needing successful delivery and action of only a single reagent (RNP) to a given cell rather than two reagents (RNP and template), 2) reduced experimental complexity, cost, and faster startup compared to those needing exogenous template, and 3) lack of artefactual insertions which may occur with exogenous templates (32). We also note that the simplicity of our system may facilitate translation to human disease in the future: *in vivo* delivery of Cas9 and mRNA (the minimally required components of our method) has already been demonstrated in the clinical trial setting (33-35). Other CRISPR-based technologies, namely base editors and prime editors, have also shown great promise in single-base editing applications. Unlike IGC, these systems can introduce a wide variety of novel base substitutions with relatively high efficiency at the user’s discretion (36). These may be the best option for base reversion in some scenarios, however, we anticipate that features of IGC may make it a preferred approach in other situations. For one, IGC does not carry the risk of editing nearby bases as base editors do, and, for another, it is a much more compact system to deliver to cells than prime editors, which require both Cas and reverse transcriptase proteins (36). Taken together, IGC is an experimentally straightforward new approach that complements existing CRISPR-based methods, and one which may have particular utility in specific scenarios, primarily when the object is to restore a disease-associated SNV to its wild-type form.

This study is, to our knowledge, the first to attempt intentional IGC of SNVs in somatic human cells, yet our results largely align with previous research that has tangentially touched on this area. A 2013 report found that a gRNA targeting a 1-bp deletion which created a novel PAM could lead to IGC in mouse zygotes (4/22 live offspring) (8), while a 2014 study in rats that used a SNV-allele-specific CRISPR KO found some of the experimentally treated zygotes had incidentally experienced spontaneous interallelic conversions (2/7 offspring) (7). In each case, there were relatively greater amounts of indels than HDR (indels in 6/22 and 5/7, respectively) (7,8). Numerous other studies have used allele-specific CRISPR in various organisms (including (8,12,37-41)), yet few have reported IGC. There are several reasons why this is not entirely surprising. First, many of these studies aimed to generate allele-specific knock-ins via exogenous HDR template, whose presence may have preempted the ability of naturally occurring IGC. Second, some of these studies targeted alleles that are substantially different than the wild-type DNA (gene fusions, larger indels), which would impede IGC. Finally, the majority of studies to date have used wild-type Cas9, while we used a high-fidelity version, and there may be differing rates of IGC based on the specificity and DNA interactions of the enzyme used. Importantly, the few studies which have described IGC following CRISPR have not targeted alleles which differ by a single SNV as we have done here, but rather targeted homologous alleles that differed by multiple SNVs or even indels (42-44). For example, while the gRNAs used in our studies differ from the non-targeted allele by just 1/20 bases (5%), previous gRNAs in previous studies have differed by 13/20 bases (65%) (42), 4/21 bases (19%) (43), or targeted an allele with a novel PAM (44).

At the same time, we acknowledge several limitations to the present study. While we attempted to test multiple different SNVs, we were limited to SNVs occurring in the biospecimens to which we had access, meaning that we were unable to ask questions such as whether SNV position within the protospacer substantially affects rates of IGC or if certain genetic loci are more/less susceptible to IGC. Moreover, we also only tested cells of hematopoietic origin, so we are unable to speak to the extent to which IGC is feasible in other somatic tissue types. And, while we find that our experiments achieved relatively more reversion to WT than introduction of indels, we were unable to entirely prevent the formation of indels in treated samples. In the patient samples, the rate of IGC was appreciable yet modest; enough to demonstrate the utility of this approach in principle but we concede further refinements may be required to maximize the utility of its application to studying disease phenotypes in primary cells. The use of electroporation would limit the utility of IGC to the *ex vivo* context, but innovations in delivery methods such as HSPC-targeted nanoparticles (45) could open up the door to *in vivo* studies.

Leading from this proof-of-concept research, there are several areas of investigation that are ripe for study. In particular, an emphasis should be placed on future studies that attempt IGC in pre-malignant cells as well as in both solid and liquid tumors. Here, we focus on somatic mutations, but there are numerous deleterious germline SNVs, including those which predispose to myeloid disease (46), which are also deserving of study. Whether it is possible to multiplex IGC and target multiple SNVs at once is also an open question. While we only test malignant cells harboring multiple genetic alterations, it would be intriguing to apply this approach to pre-cancerous cells with a clonal mutation (e.g., clonal hematopoiesis), where a single SNV might be the only feature defining a clone. But even as we describe it here, we think that IGC holds promise as a straightforward tool to study the potential benefit that may come from clonal reductional of a harmful SNV. Similarly, it could be applied to testing whether specific SNVs contribute to therapeutic resistance, by decreasing (or increasing) the SNV VAF prior to drug testing. The workflow we describe is highly analogous to a straightforward CRISPR KO protocol, and the only essential components for IGC are the high-fidelity Cas enzyme and gRNA, which may be readily obtained from a variety of scientific vendors. The ability to reduce specific SNVs while leaving wild-type DNA intact could have significant positive implications for the study of disease and pre-disease states.

In conclusion, SNV-specific interallelic gene conversion is a simple CRISPR-based approach that can revert a heterozygous SNV allele to its counterpart allele which we have demonstrated works across multiple contexts in leukemic cells. We anticipate that this technique may find broad application in basic and translational medical research focused on the contribution of SNVs to disease pathogenesis.

## AUTHOR CONTRIBUTIONS

*Conceptualization*: A.S. & M.S. *Formal analysis*: A.S., H.R., S.O., & M.S. *Funding acquisition*: M.S. *Investigation*: A.S., D.B., H.R., A.G., S.O., J.W., & L.F *Methodology*: A.S., H.R., & M.S *Validation*: A.S., D.B., S.O. & H.R. *Supervision*: M.S. *Visualization*: A.S. *Writing – original draft*: A.S. & M.S. *Writing – review & editing*: A.S., D.B., H.R., A.G., S.O., J.W., L.F., & M.S.

## ACKNOWLEDGEMENTS

A.S. received financial support from the US National Institutes of Health (NIH) under Ruth L. Kirschstein National Research Service Award F30DK127699 from the NIDDK, and T32GM007347 from the NIGMS. M.S. receives funding from Leukemia and Lymphoma Society, the E.P. Evans Foundation, The Biff Ruttenberg Foundation, the Adventure Alle Fund, the Beverly and George Rawlings Directorship, and the NIH grants: 1R01CA262287, 1U01OH012271, P30 CA068485.

Flow cytometry was conducted in the VUMC Flow Cytometry Shared Resource, which is supported by the Vanderbilt Ingram Cancer Center (P30 CA68485) and the Vanderbilt Digestive Disease Research Center (DK058404). The content is solely the responsibility of the authors and does not necessarily represent the official views of the National Institutes of Health.

