## Supplementary Information for "Repair of leukemia-associated single nucleotide variants via interallelic gene conversion"

**SUPPLEMENTARY MATERIAL**
**Supplementary Tables: Page 2-3**
**Supplementary Figures: Pages 4-5**

**Supplementary Table 1. PCR primers used for Sanger sequencing.**

| Locus | Forward | Reverse |
|---|---|---|
| ***DNMT3A* p.R882** | GAGGAGTTGGTGGGTGTGAG | GGGTTCTGTTTTGCGTGACC |
| ***NRAS* p.G12** | CCTTAGGTGCCACTTACTGAA | GGCCGATATTAATCCGGTGT |
| ***ASXL1* p.Y591** | CTTCTCTGAATGGTGTGTTAT | GCCAGTTCCTTTCTCTAATGT |

**Supplementary Table S2. Phased indels in primary acute myeloid leukemia samples.**

| Sample | gRNA | Replicate | Total Reads | Indels on Wild-type Alleles (%) | Indels on Mutant Alleles (%) | Indels Unable to be Phased (%) |
|---|---|---|---|---|---|---|
| AML 01 | Scramble | 1 | 1207 | 1 (0.1%) | 0 (0%) | 0 (0%) |
| | | 2 | 2102 | 0 (0%) | 0 (0%) | 0 (0%) |
| | | 3 | 279 | 0 (0%) | 0 (0%) | 0 (0%) |
| | Q748X | 1 | 1334 | 0 (0%) | 0 (0%) | 0 (0%) |
| | | 2 | 870 | 2 (0.2%) | 1 (0.1%) | 5 (0.6%) |
| | | 3 | 1125 | 2 (0.2%) | 0 (0%) | 0 (0%) |
| AML 02 | Scramble | 1 | 2026 | 0 (0%) | 0 (0%) | 0 (0%) |
| | | 2 | 1805 | 0 (0%) | 0 (0%) | 0 (0%) |
| | | 3 | 1716 | 0 (0%) | 0 (0%) | 0 (0%) |
| | R882H | 1 | 850 | 1 (0.1%) | 0 (0%) | 0 (0%) |
| | | 2 | 1082 | 0 (0%) | 0 (0%) | 0 (0%) |
| | | 3 | 1363 | 2 (0.1%) | 0 (0%) | 0 (0%) |

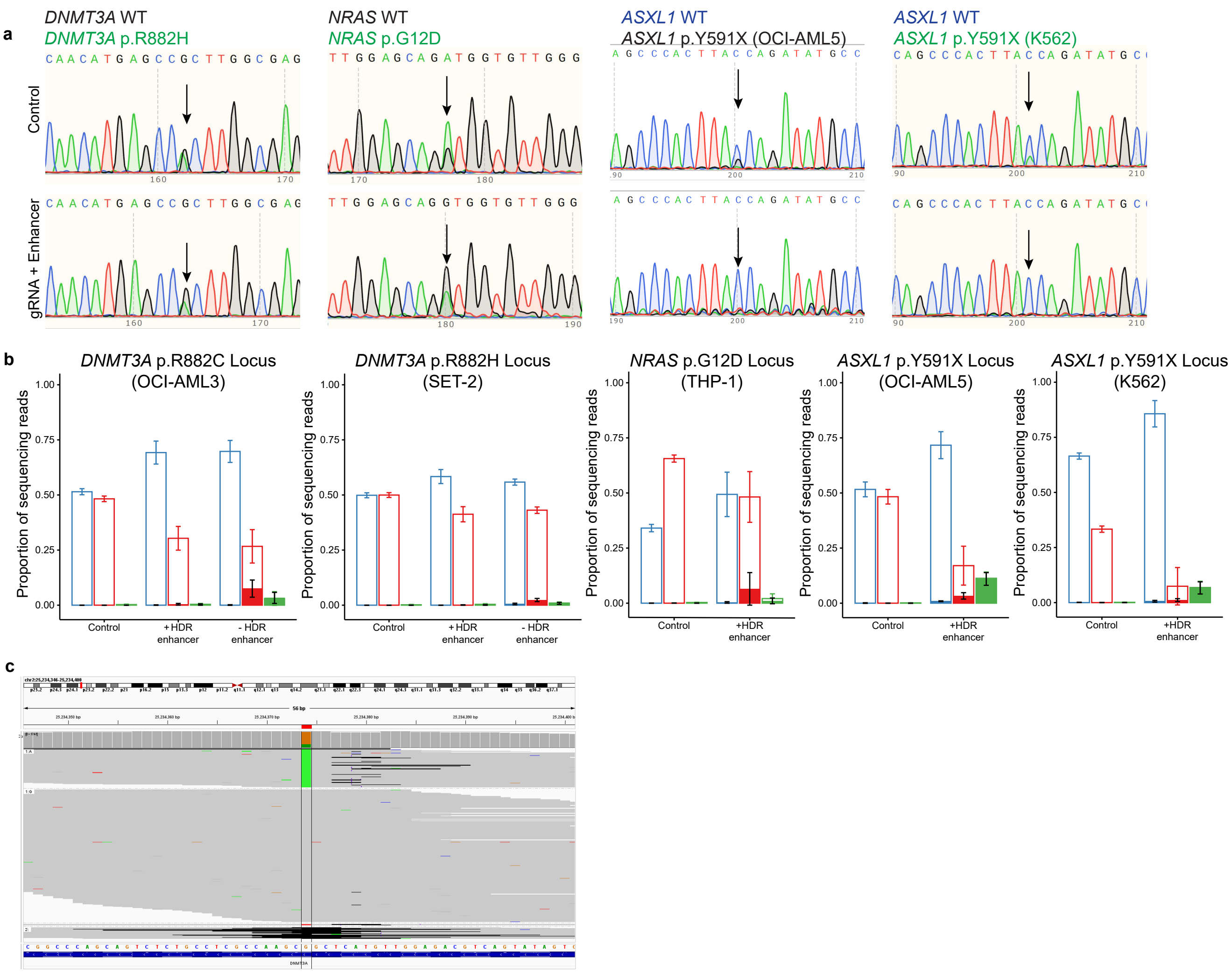


**Supplementary Figure S1. Additional description of cell line samples following gene editing.** (A) Representative Sanger traces for SET-2, THP-1, OCI-AML5, and K562 cells. (B) Proportion of all wild-type (blue), mutant (red), or unphased reads (green; empty bars) and the proportion of indel-containing wild-type, mutant, or unphased reads (shaded bars) for OCI-AML3, SET-2, THP-1, OCI-AML5, and K562 cells. Bar plots and error bars represent mean and standard deviation. (C) Integrated Genome Viewer locus plot depicting indels in downsampled NGS reads at the DNMT3A p.R882C locus following OCI-AML3 treatment with mutant-specific gRNA without homology-directed repair (HDR) enhancer. The targeted single nucleotide variant is bordered by black vertical lines, with mutant allele in green, wild type allele in gray, and deletions shown as black horizonal lines.

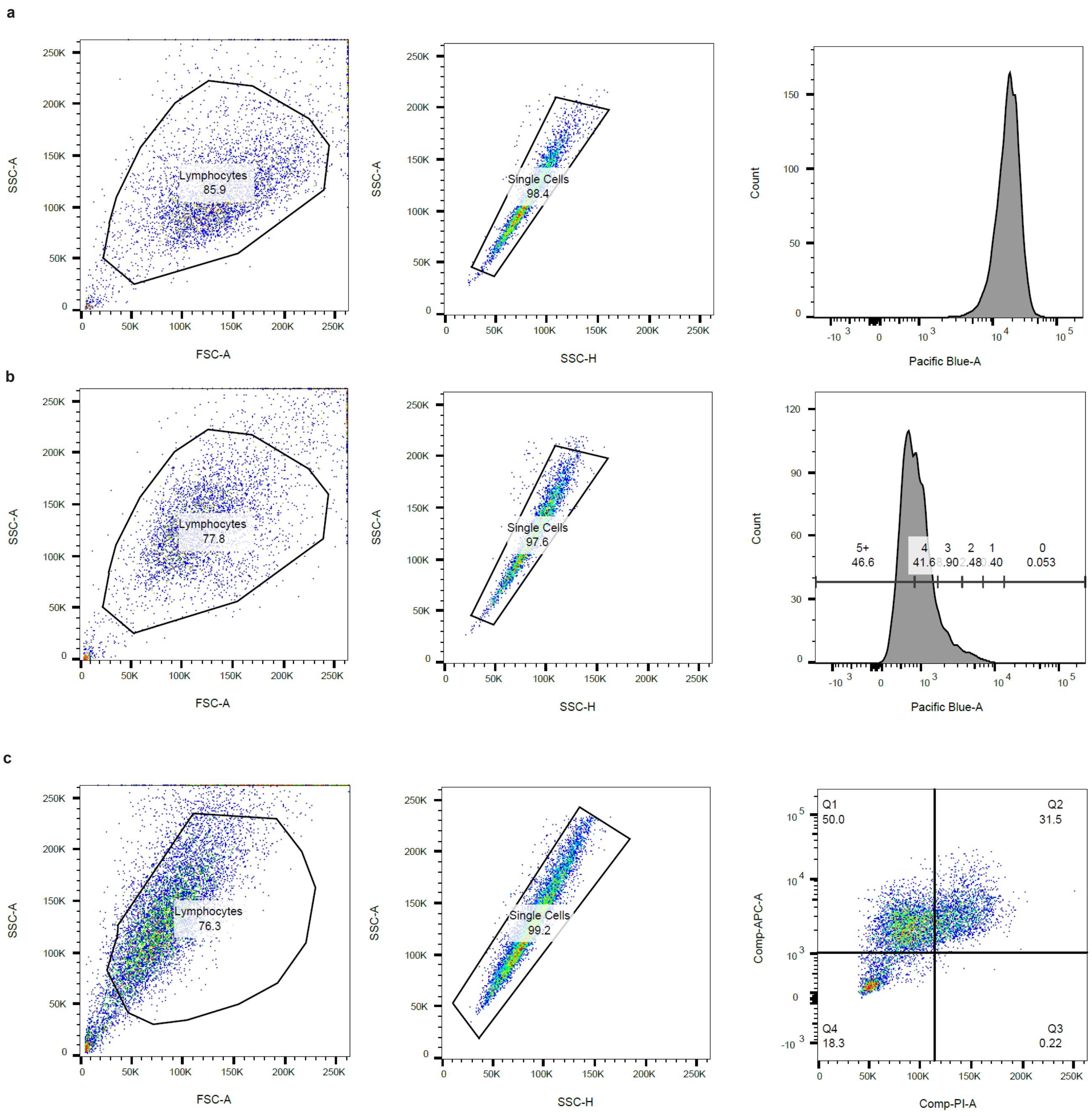


**Supplementary Figure S2. Flow cytometry gating strategies.** FACS gating strategy for CellTrace assay, showing (A) a sample fixed at time = 0 and (B) fixed at time = 72 hours. (C) FACS gating strategy for Ki-67 cell cycle assay.
